# Inference of evolutionary forces acting on human biological pathways

**DOI:** 10.1101/009928

**Authors:** J.T. Daub, I. Dupanloup, M. Robinson-Rechavi, L. Excoffier

## Abstract

Because natural selection is likely to act on multiple genes underlying a given phenotypic trait, we study here the potential effect of ongoing and past selection on the genetic diversity of human biological pathways. We first show that genes included in gene sets are generally under stronger selective constraints than other genes and that their evolutionary response is correlated. We then introduce a new procedure to detect selection at the pathway level based on a decomposition of the classical McDonald-Kreitman test extended to multiple genes. This new test, called 2DNS, detects outlier gene sets and takes into account past demographic effects as well as evolutionary constraints specific to gene sets. Selective forces acting on gene sets can be easily identified by a mere visual inspection of the position of the gene sets relative to their 2D null distribution. We thus find several outlier gene sets that show signals of positive, balancing, or purifying selection, but also others showing an ancient relaxation of selective constraints. The principle of the 2DNS test can also be applied to other genomic contrasts. For instance, the comparison of patterns of polymorphisms private to African and non-African populations reveals that most pathways show a higher proportion of non-synonymous mutations in non-Africans than in Africans, potentially due to different demographic histories and selective pressures.

## Introduction

In the search for genomic signals of natural selection (e.g. reviewed in Nielsen 2005), there has been a recent shift from single gene to gene set approaches, where the focus moved to gene networks, pathways and interacting genes (Daub et al. 2013; Fraser 2013; Zhang G et al. 2013; Berg and Coop 2014). Studying groups of functionally related genes makes biological sense for several reasons. First, as selection is acting on phenotypic traits usually controlled by many genes (Stranger et al. 2011), we would expect it to affect multiple genes or a whole pathway rather than a single gene. A biological pathway or a gene network is therefore a more natural unit for selection tests. Second, mutations in one gene can induce the modification of functionally connected genes in order to adapt to or compensate for the initial change (Pazos and Valencia 2008), and such co-evolution can lead to a cascade of evolutionary changes within a gene network. Third, many small effect mutations can together have a large effect on polygenic traits. It has been suggested that selection on such traits, termed polygenic selection, usually acts on standing variation at several loci at the same time. In the long run this would lead to an increase in the frequency of a suitable combination of alleles that control the favored phenotype (Pritchard et al. 2010).

For example, using levels of population differentiation between human groups, we have recently found several pathways involved in immune response to have been under positive selection in recent human history (Daub et al. 2013). In the current paper we investigate if and what type of polygenic selection could have acted at different stages of human evolution, by comparing patterns of diversity between humans and chimpanzees. The rationale is that since selective pressures could have changed over time for some biological pathways, recent or old episodes of selection cannot be detected by only looking at current human diversity. Our method contrasts patterns of fixed and polymorphic mutations in humans, allowing us to detect various selective pressures having affected our species at different periods of its evolution.

Our approach is based on statistics previously used in the classical McDonald-Kreitman (MDK) test for positive selection (McDonald and Kreitman 1991), a test that compares the ratio of the number of non-synonymous substitutions to synonymous substitutions between species (DN/DS) to the same ratio within species (PN/PS). Assuming that synonymous mutations are neutral, a higher DN/DS than PN/PS ratio is expected in case of ancient adaptive selection, because positively selected mutations rising to fixation in a population would contribute more to divergence than to polymorphism. The related statistic *α* = 1-(*DS*×*PN*)/ (*DN*×*PS*)has been used to estimate the proportion (*α*) of adaptive substitutions in a genome (Smith and Eyre-Walker 2002). A limitation is that the same *α* value can be obtained for different combinations of PN/PS and DN/DS ratios. In order to account for this problem, we introduce here a testing procedure, based on a two-dimensional decomposition of *α*, which identifies gene sets departing from a genome-wide null distribution, and leads to a better interpretation of the results by a visual inspection of bivariate distributions.

## Material and Methods

### Data handling and collection

#### Ensembl gene data

We downloaded the exon coordinates of protein coding human genes from Ensembl (Ensembl version 64, September 2011, Flicek et al. 2014). We then only considered those genes (and their corresponding coding exons) that have a chimp ortholog with a ‘known’ status as defined in Ensembl version 64, and genes having a chimp ortholog without ‘known’ status, but with a ‘known’ mouse ortholog. The second group represents genes that most probably have a real ortholog in chimp, but are not annotated as such (yet) due to the lower quality of the chimp sequence. This left us with 18,078 genes and we will refer to this set as *G*_*Ensembl*_. We computed the total exon length of each gene by summing the length of all corresponding exons, but only counted a site once if it was part of two overlapping exons.

#### Human SNPs

Human single nucleotide polymorphisms (SNPs) were inferred from the comparison of the whole genomes of 42 unrelated individuals sequenced by Complete Genomics at a depth of 51-89X coverage per genome (Drmanac et al. 2010). The 42 individuals were sampled in 3 African populations (4 Luhya from Webuye, Kenya; 4 Maasai from Kinyawa, Kenya; 9 Yoruba from Ibadan, Nigeria) and 5 non-African populations (9 Utah residents with Northern and Western European ancestry from the CEPH collection; 4 Han Chinese from Beijing; 4 Gujarati Indians from Houston, Texas, USA; 4 Japanese from Tokyo; 4 Toscans from Italy). The SNPs were divided into three categories: sites which are polymorphic in Africans only (African SNPs), sites polymorphic in non-Africans only (non-African SNPs) and sites polymorphic in both Africans and non-Africans (shared SNPs). The shared SNPs presumably arose before the migration of modern humans out of Africa and are therefore depleted from recent deleterious mutations that otherwise could distort selective signals. This group of SNPs was used in the 2DNS test (Figure 1 and Figure S1). The African and non-African SNPs were used in a further analysis to compare demographic patterns between the two regions (Figure 2).

**Figure 1.**
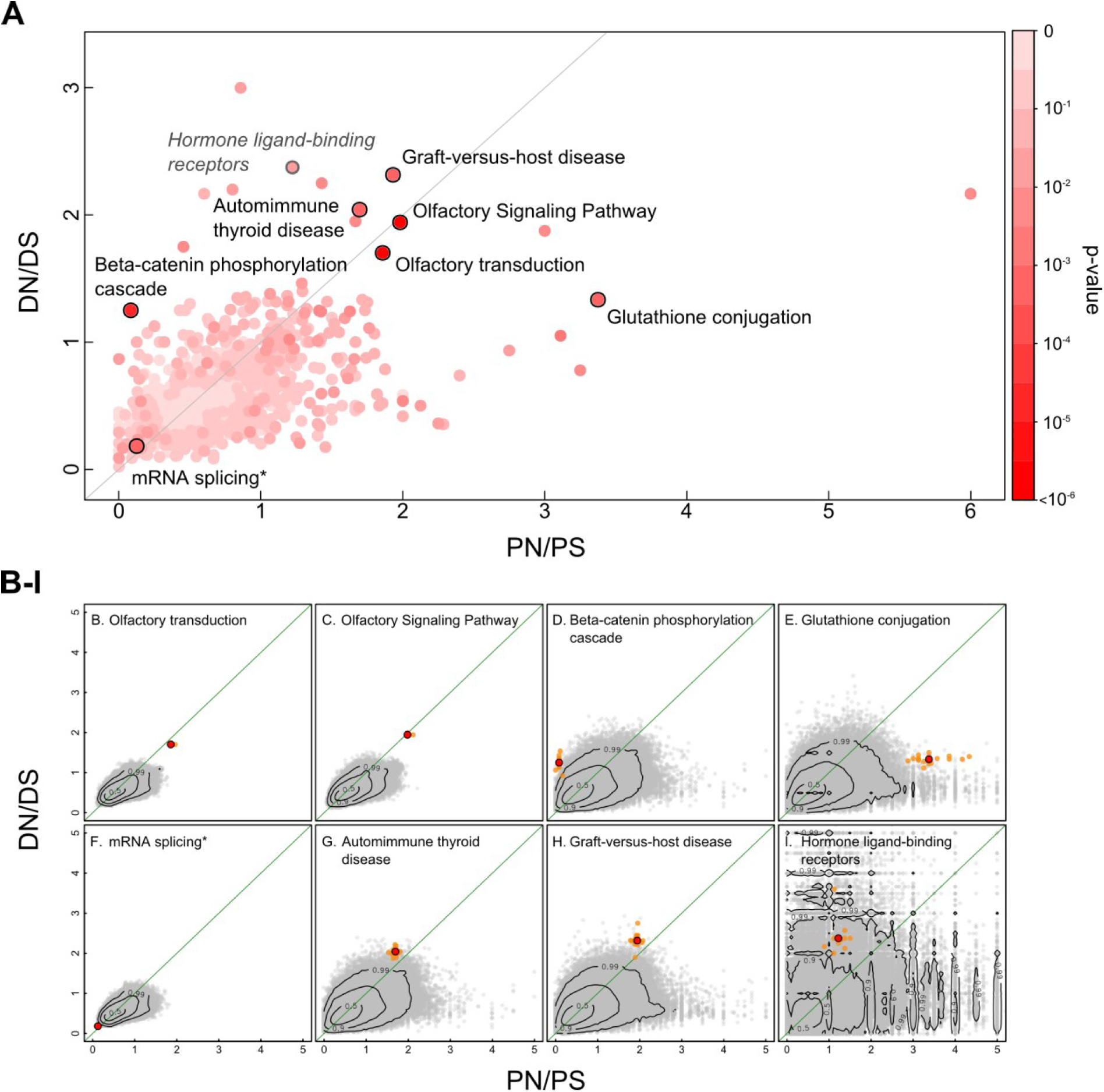
Results from the 2DNS test. Here, shared SNPs (SNPs that are polymorphic both in African and non-African populations) are compared to fixed substitutions in the human branch. (A): Distribution of DN/DS and PN/PS ratios of all tested pathways. Each dot represents a pathway with color corresponding to its significance. The seven pathways that scored significant (q-value < 0.1; Table 2) in the 2D tests are highlighted by a black circle. Note that seemingly outlier gene sets may not reach significance due to their small size, which increases the variance of their null distribution. For example, the *Hormone ligand-binding receptors* pathway (depicted in grey; panel I), has a high DN/DS ratio, but because of its small size (10 genes, 24 kb total exon length) its null distribution is very widespread in the 2D space. (B-H): Null distributions for seven significant (q<0.1) pathways; (I): null distribution for the *Hormone ligand-binding receptors* pathway as a typical example of a non-significant outlier. The observed positions of gene sets are indicated as red dots in the DN/DS-PN/PS plane, whereas the empirical null distribution is shown as grey dots. Orange dots show the scores of the jackknifed gene sets. The contour lines mark the proportion (0.5, 0.9, and 0.99) of the null distribution that falls within these areas.

**Figure 2.**
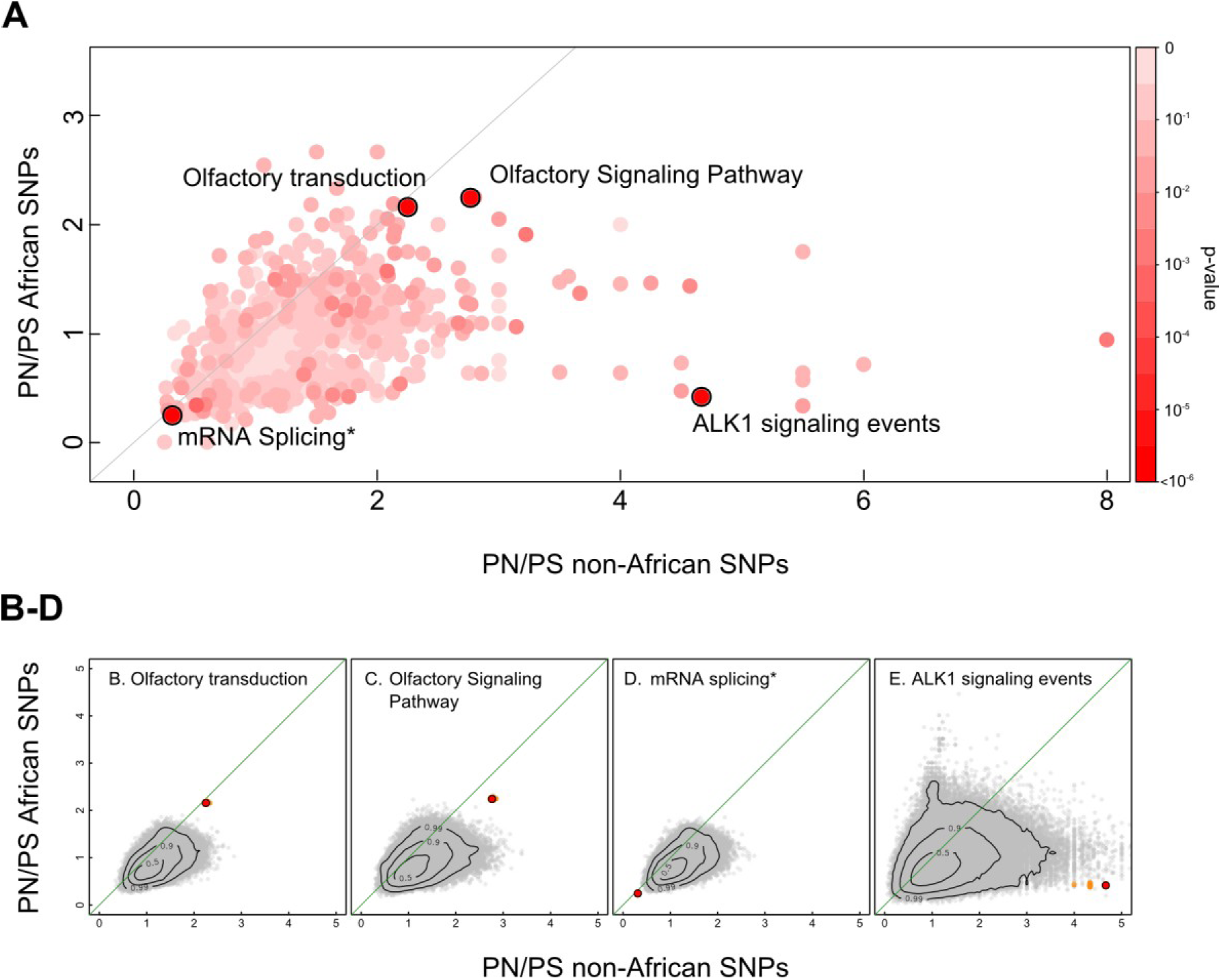
Results of the 2DNS test on PN/PS ratios of all tested pathways. Here, non-African SNPs (SNPs that are only polymorphic in non-African populations) are compared to African SNPs (SNPs unique to African populations). (A): Distribution of DN/DS and PN/PS ratios of all tested pathways. Each dot represents a pathway with color corresponding to its significance. The five pathways that scored significant (q-value < 0.1) in the 2DNS tests are highlighted by a black circle. (B-D): Null distributions for four significant (q<0.1) pathways. The observed positions of gene sets are indicated as red dots in the DN/DS-PN/PS plane, whereas the empirical null distribution is shown as grey dots. Orange dots show the scores of the jackknifed gene sets. The contour lines mark the proportion (0.5, 0.9, and 0.99) of the null distribution that falls within these areas.

#### Human Chimp substitutions

Substitutions between human and chimp were inferred from the comparison between the reference genomes of the two species, using the syntenic net alignments between hg19 and panTro2 available on the UCSC platform (Karolchik et al. 2012). The inferred mutations were then placed on the human-specific or chimp-specific branches of the tree by comparing the nucleotides observed in human and chimp to the orthologous base in the orang genome (Figure S1), which was obtained using the syntenic net alignments between hg19 and ponAbe2 available on the UCSC platform (Karolchik et al. 2012); only mutations which could be placed unambiguously on the human or chimp branch were used.

#### Annotating synonymous and non-synonymous substitutions

We mapped the mutations to the coding exons of the human genes in the *G*_*Ensembl*_ set. We classified the mutations as either synonymous or non-synonymous using ANNOVAR (Wang et al. 2010). Our polymorphism and divergence data cover a largely overlapping part of the exome: 89.2% of the exome is fully sequenced in all 42 Complete Genomics samples, whereas the human-chimp alignment covers 96.3% of the exome, resulting in 86.8% of the exome covered by both data sets.

#### Ensembl to Entrez Gene ID conversion

Since genes in Biosystems gene sets are annotated with Entrez gene IDs, we mapped the Ensembl gene IDs in *G*_*Ensembl*_ to Entrez IDs by constructing a one-to-one Ensembl - Entrez conversion table. To do this, we first downloaded from the NCBI Entrez Gene website (Maglott et al. 2011) a gene list (*G*_*Entrez*_) with 19,759 ‘current’ protein coding human genes (http://www.ncbi.nlm.nih.gov/gene, accessed on February 7, 2013). Next, we collected conversion tables (containing one-to-many or many-many mappings) from three sources: Ensembl (version 64, http://sep2011.archive.ensembl.org/biomart), NCBI (ftp://ftp.ncbi.nih.gov/gene/DATA, accessed on February 7, 2013), and HGNC (http://www.genenames.org/biomart/, accessed on September 3, 2012). From these tables we only kept rows with genes in *G*_*Ensembl*_ and *G*_*Entrez*_. We then pooled the three tables and counted the occurrences of each unique Ensembl ID - Entrez ID combination. For each Ensembl ID we kept the Ensembl - Entrez match with the highest count (in case of multiple options we took the first one) and repeated this for each Entrez ID. The resulting table contained 17,474 one-to-one Ensembl to Entrez gene ID conversions and we used the genes from this table (gene list *G*) for our further analyses.

#### Gene sets

We downloaded 2,402 human gene sets from the NCBI Biosystems database (Geer et al. 2010) (http://www.ncbi.nlm.nih.gov/biosystems, accessed on February 2, 2013). We excluded genes that could not be mapped to the gene list *G* and removed gene sets with less than 10 genes. Furthermore, we identified 95 groups of (nearly) identical gene sets, namely sets sharing at least 95% of their genes, and replaced each group by a union gene set (in the text their name is followed by ‘*’) consisting of all genes in that group. The remaining collection of 1,366 gene sets (*S*) was used for our analyses. Note that, for ease of reading, we use both the terms pathway and gene set interchangeably throughout the main text; strictly speaking only a small proportion (<2%) of the sets in *S* are labeled as ‘complex’ rather than ‘pathway’ in the Biosystems database.

### Data analysis

#### Test of evolutionary similarities between genes of a given gene set

To assess whether genes in sets tend to share the same evolutionary properties, we tested for significant differences in gene-level DN/DS (or PN/PS) ratios between gene sets. First, we estimated the variance component due to differences among gene sets 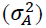 for all 1,366 sets using an analysis of variance (ANOVA) framework (See Supplemental Text S1). We constructed the expected null distribution of 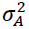 under the hypothesis that there are no differences between groups, by repeatedly (*N*=10,000) recalculating 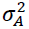 after permuting DN/DS and PN/PS ratios among genes. Next, to assess that the significance was not due to one or a few outlier gene sets, we repeatedly (*N*=10,000) sampled randomly 100 sets from the list of 1,366 gene sets and computed 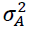 on these 100 gene sets only. We built the expected null distribution by performing the same sample procedure, but permuting DN/DS and PN/PS ratios among genes for each drawing of 100 genes sets. (See also Figure S2). Note that the fact that genes can occur in multiple gene sets has no impact on the results, as we keep the gene set definitions fixed when creating the null distribution. Therefore the amount of overlap between gene sets is the same in both null and observed distributions.

Note that we excluded 5,314 (3,049) genes that had zero DS (PS), leading to undefined N/S ratios, when comparing DN/DS (or PN/PS) ratios between genes (as shown in Figure S3 and in the ANOVA test described above). In the other analyses, we could use all 17,474 genes of our gene list *G* to compute DN/DS (or PN/PS) ratios at the gene set level.

#### Gene and gene set level DN/DS and PN/PS ratios

To calculate the DN/DS ratio for a gene set, we summed up separately the non-synonymous and synonymous fixed mutations found in all genes belonging to a given gene set (Figure S1) and took their ratio as

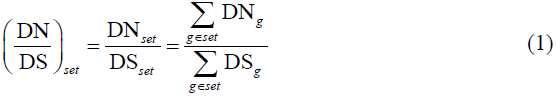

The PN/PS ratio was calculated in a similar way. In a few cases where the same mutation occurred in more than one gene of the same gene set (due to gene overlap or exon sharing, around 0.1% of the mutations), it was counted only once.

### Null distributions

#### Gene set properties taken into account when building empirical null distributions

We constructed empirical null distributions of N/S ratios by creating a series of random gene sets and recomputing new statistics on these gene sets. As genes in gene sets might have specific evolutionary properties, randomly sampling genes from the gene list *G* would not create a representative null distribution. Instead, we took into account the negative correlation found between the number of gene sets a given gene belongs to, and the gene N/S ratio. We also took into account the fact that genes belonging to the same pathway tend to share the same evolutionary regime (Figure S2). To preserve the evolutionary properties of gene sets, we applied a bootstrap sampling method, where we sampled genes from unions of gene sets that share a given proportion of their genes (see next section).

In addition, we also considered the fact that gene sets with a large ‘set length’ (defined as the total exon length of its genes), accumulate on average more mutations and for this reason show a lower variance in N/S rates. We therefore created null distributions for different set length categories and tested each pathway against the appropriate null distribution.

#### Construction of empirical bootstrap null distributions

In short, we first created random gene sets of different sizes (number of genes), where we sampled the size according to the real size distribution and next computed the set length of the random sets and assigned them to different set length bins.

We created a random gene set as follows. First, we drew a gene set size *k* from the geometric distribution that approximated the real gene set size distribution, but allowed for sampling gene sets smaller than 10 (the minimum gene set size) in order to obtain sufficient sets to fill the smallest set length bins:

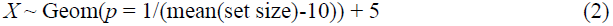

Next, we sampled a gene set from the collection *S* of all gene sets. For each sampled gene set *S′*, we collected all genes that are the union of genes in *S′* and genes in any set *S″* that overlaps with *S′*, in the sense that their Jaccard similarity coefficient (defined as [S′ ∩ S″]/[S′ U S″]) is larger than 0.25. If the union contains less than min(250, *k*) genes, it was discarded, otherwise we randomly sampled *k* genes (without replacement if *k*<250, otherwise with replacement) from this union to form a random gene set. We repeated this 75 × *N* (*N*=400,000) times. For each observed and random gene set we calculated its set length and distributed the sets according to their set length over 75 equally sized bins, explaining the 75 factor in the previous sentence.

In very few occasions the N/S ratio of a (random) gene set was not defined, because S was zero. In these cases we removed the gene set from the null distribution or from the group of sets that were tested. To still have bins of size *N*, we therefore actually created 75 × *N* × 1.1 gene sets, removed the sets that did not have a defined N/S ratio and took the first *N* remaining gene sets for each bin.

#### Alternative null distributions without gene set properties

To investigate the effect of not taking into account some or all of the properties of gene sets when creating a null distribution, we used several alternative sampling schemes consisting of: (1) randomly sampling from the whole gene list *G*; (2) sampling from all genes that are part of at least one gene set; and (3) sampling genes with a probability proportional to the number of gene set a gene belongs to. Similar to the empirical bootstrap null distribution described above, we corrected in all cases for gene set length, by creating null distributions for specific set lengths and testing a pathway against the corresponding null distribution. Results based on these alternative null distributions are shown in Supplemental Figure S4.

### Tests of selection

#### A new test of polygenic selection (2DNS test)

We developed a new test for polygenic selection (named 2DNS), which can be regarded as a decomposition of the constitutive elements of a MDK test, DN/DS and PN/PS. Our 2DNS test aims at detecting gene sets that are outliers in the two dimensional DN/DS - PN/PS plane relative to an empirical null distribution of joint DN/DS and PN/PS ratios. The p-value of a gene set is estimated from the joint N/S density on a two dimensional grid. We first used the R function *kde2d* (taking bandwidth = 0.15, maximum grid size = 10 × 10 and grid point distance = 0.05) to estimate the probability density of each of the grid points. We then bilinearly interpolated the N/S coordinates of the observed gene set on the grid of the null distribution and the corresponding density with the R function *interp.surface*. The p-value of the gene set is finally calculated by integrating over the densities of grid points that have the same or lower density as the density of the observed gene set (Figure S5). Note that because we use a kernel density estimation, p-values much smaller than the inverse of number of elements in the null distribution can be obtained.

#### Correction for multiple tests

We FDR corrected the p-values from our tests by calculating the q-value of each gene set using the function *qvalue* from the R package *qvalue* with default settings. We retained those gene sets with a q-value below 0.1, meaning that we expect 10% of our final candidates to be false positives.

#### Jackknife approach to detect pathways with outlier genes

We used a jackknife approach to examine the effect of individual genes of a given gene set on our results. For each significant pathway, we repeatedly removed one gene and recalculated the DN/DS and PN/PS values. These jackknife scores are depicted in Figure 1B-I. Pathways where one of the jackknife scores resulted in a much higher p-value were probably scoring significant because one gene has extreme values and such pathways were not considered as candidate gene sets for *polygenic* selection

#### Classical MDK test at gene set level

The classical MDK test for positive selection tests at the gene level whether the DN/DS ratio is significantly larger than the PN/PS ratio. We extended this procedure to the gene set level by creating for each gene set a 2 × 2 contingency table containing its DN_*set*_, DS_*set*_, PN_*set*_ and PS_*set*_ counts as defined in eq. (1) and testing whether the odds ratio of the table (OR = DS·PN/DN·PS) is significantly deviating from (smaller than) one with a two-sided (one-sided) Fisher exact test, using the R function *fisher.test*.

#### Comparing gene set level alpha values with an empirical null distribution

We calculated a gene set analog of *α* for each set, here defined as

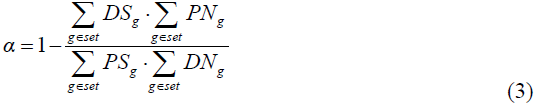

and compared it to the *α* values of an empirical null distribution with corresponding set length.

## Results

### Genes in biological pathways are conserved and share evolutionary properties

We first investigated whether genes in pathways have different evolutionary properties than genes that are not part of a pathway. We downloaded a collection of 1,366 pathways from the Biosystems database (Geer et al. 2010) and contrasted the genetic diversity of genes that occur in one or more gene sets with genes that are not part of any gene set. We used single nucleotide polymorphisms (SNPs) inferred from the Complete Genomics collection of 42 human individuals from three African and five non-African populations (Drmanac et al. 2010), as well as the fixed differences between humans and chimpanzees that were assigned to the human branch. Interestingly, we find that the non-synonymous to synonymous ratios are lower for genes in gene sets than for genes that are not part of a gene set, both for mutations within human populations (PN/PS) and for fixed mutations on the human lineage (DN/DS). It suggests that genes belonging to gene sets are globally under more severe evolutionary constraints (see Table 1).

**Table 1.**
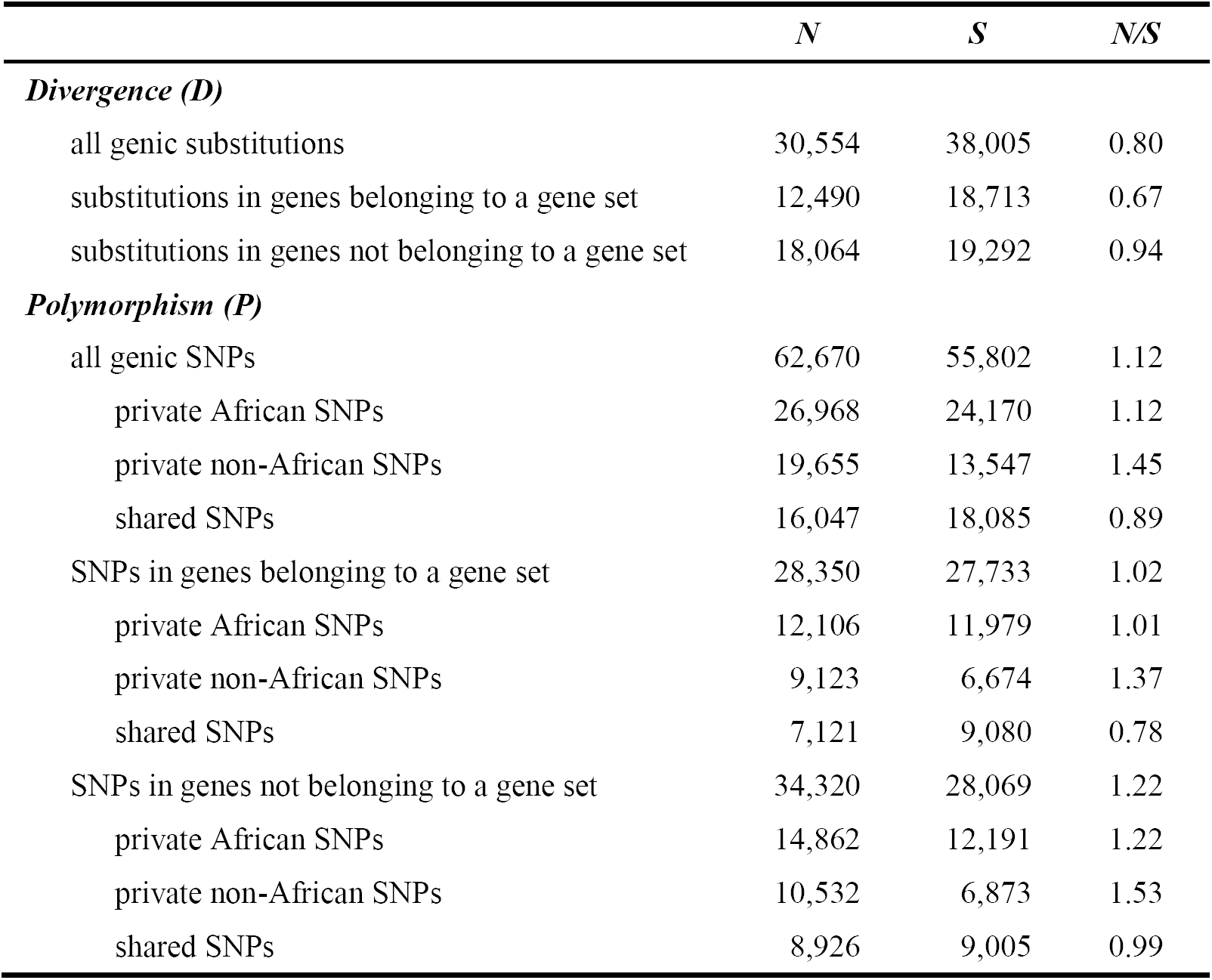
Counts and ratios of non-synonymous (*N*) and synonymous (*S*) fixed mutations and polymorphisms in the human lineage.

In addition, we find that genes that are part of many pathways (10 or more) are even more constrained than genes occurring in a single or a few pathways, as they show significantly lower DN/DS and PN/PS ratios (p<1e-6, Mann-Whitney test, Figure S3). Overall, there is a small but significant negative correlation between the DN/DS ratio of a gene and the number of pathways it belongs to (Pearson’ s r = -0.13, p<2.2e-16). These results make sense, since genes that are part of many pathways often have an essential role or have several functions, suggesting a potential pleiotropic effect of mutations in these genes, and thus a higher chance for them to be deleterious. These findings are also consistent with earlier reports showing that highly connected proteins are usually more essential (Jeong et al. 2001; Wuchty 2004) or evolving at a slower rate (Fraser et al. 2002; Saeed and Deane 2006). These results further imply that tests of selection bearing on single genes should be performed separately for genes not part of gene sets, or that their lower levels of evolutionary constraints be taken into account in future testing procedures to minimize false positives.

Interestingly, we observe that gene-specific DN/DS and PN/PS ratios are more similar within than between gene sets, and that the variance in N/S ratios due to differences between gene sets represents around 5% (Figure S2) of the total variance in N/S ratios (non-parametric ANOVA test, p-value<1e-4; see Material and Methods and Text S1). This variance component also represents the average correlation between N/S ratios of two genes belonging to the same gene set relative to two genes from different gene sets. This positive correlation of about 5% suggests that genes in a pathway share evolutionary properties and have a correlated evolutionary response (i.e. most are rather conserved or most are under weak selection). These results are in keeping with the observation of Fraser et al. (2002) that interacting proteins have similar evolutionary rates, which the authors explained by the occurrence of compensatory changes in interacting proteins.

### Measuring and testing selective constraints at the pathway level

We are interested here in finding a way to determine the extent and the type of selection acting in gene sets. Conventional McDonald-Kreitman tests such as those based on the *α* statistic are usually used to detect only positive selection (but note that a negative *α* is an indicator of purifying selection). Furthermore, *α* is based on a ratio of ratios, and a given high *α* value, taken to be indicative of positive selection, can be obtained with a high DN/DS or a low PN/PS, which can be due to different evolutionary forces. To address these issues, we propose here a new test of polygenic selection that is more informative about the respective importance of DN/DS and PN/PS ratios, and that detects outlier pathways for different types of selection. Our aim with this test, referred to hereafter as 2DNS test, is to find gene sets that have evolved differently than other gene sets, in that they have unusual DN/DS or PN/PS combinations compared to other sets. First, we sum up the PN, PS, DN and DS counts over all genes in a gene set to obtain gene set level DN/DS and PN/PS ratios (as shown in eq. (1)). Second, we create genome-wide empirical null distributions of random gene sets that take into account gene set size, as well as potentially shared evolutionary properties of gene sets. Note that the use of these null distributions also allows us to control for the past demographic history shared among all gene sets, and for the overall selective constraint acting on coding regions. Third, the joint probability density distribution of DN/DS vs. PN/PS ratios is estimated on a two-dimensional grid directly from the empirical null distribution. The p-value for each gene set is finally obtained by integrating the joint density of all grid points that have a similar or lower density than the gene set of interest defined by its ‘N/S coordinates’ (see Figure S5).

We compared the distribution of gene set p-values with a uniform distribution on a QQ plot to confirm that our testing procedure was well behaved (Figure S4D). The fact that the observed p-values are close to the expected uniform distribution suggests that our null distribution correctly represents the properties of the gene sets. Note that naively constructed null distributions that ignored some or all of the evolutionary properties of gene sets led to strongly underestimated p-values which would have translated in a large number of false positive gene sets (see Figure S4A-C).

We applied the 2DNS test to the comparison between fixed mutations on the human branch and polymorphisms that are found both in African and non-African populations. These ‘shared’ polymorphisms come from relatively ancient mutations that predated the migration out of Africa, and we thus expect that slightly deleterious mutations – which can distort selective signals (Eyre-Walker 2002; Fay et al. 2002; Parsch et al. 2009) – have been purged from this group, which is confirmed by Table 1.

The 2DNS test identifies 7 gene sets with a q-value < 0.1 (Figure 1 and Table 2). These gene sets are located well outside the bulk of the 2D null distributions in different directions (Figure 1B-1H), suggesting that they are subject to different forms of selection. For each of the significant pathways, we repeatedly recalculated the DN/DS and PN/PS values leaving one gene out in turn (‘jackknifing’) to examine to which extent the score depends on single genes (Figure 1B-1I).

**Table 2.**
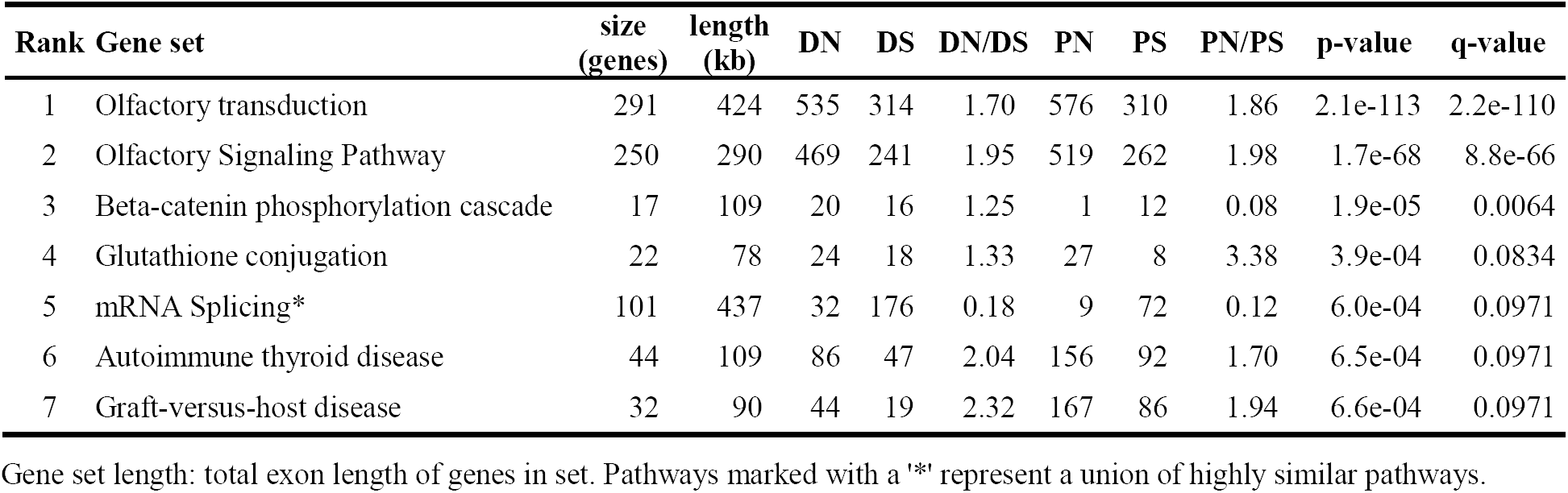
Pathways scoring a q-value < 0.1 in the 2DNS test comparing fixed mutations in the human branch (DN/DS) and polymorphisms shared between African and non-African populations (PN/PS).

| Rank | Gene set | size<br>(genes) | length<br>(kb) | DN | DS | DN/DS | PN | PS | PN/PS | p-value | q-value |
| --- | --- | --- | --- | --- | --- | --- | --- | --- | --- | --- | --- |
| 1 | Olfactory transduction | 291 | 424 | 535 | 314 | 1.70 | 576 | 310 | 1.86 | 2.1e-113 | 2.2e-110 |
| 2 | Olfactory Signaling Pathway | 250 | 290 | 469 | 241 | 1.95 | 519 | 262 | 1.98 | 1.7e-68 | 8.8e-66 |
| 3 | Beta-catenin phosphorylation cascade | 17 | 109 | 20 | 16 | 1.25 | 1 | 12 | 0.08 | 1.9e-05 | 0.0064 |
| 4 | Glutathione conjugation | 22 | 78 | 24 | 18 | 1.33 | 27 | 8 | 3.38 | 3.9e-04 | 0.0834 |
| 5 | mRNA Splicing* | 101 | 437 | 32 | 176 | 0.18 | 9 | 72 | 0.12 | 6.0e-04 | 0.0971 |
| 6 | Autoimmune thyroid disease | 44 | 109 | 86 | 47 | 2.04 | 156 | 92 | 1.70 | 6.5e-04 | 0.0971 |
| 7 | Graft-versus-host disease | 32 | 90 | 44 | 19 | 2.32 | 167 | 86 | 1.94 | 6.6e-04 | 0.0971 |

Gene set length: total exon length of genes in set. Pathways marked with a '\*' represent a union of highly similar pathways.

### Significant pathways have been influenced by different types of selection

The two highest scoring sets, *Olfactory transduction* and *Olfactory Signaling Pathway*, have both high DN/DS and high PN/PS ratios (Figure 1B-C), in line with their known relaxed constraints in primates, in particular in humans (Gilad et al. 2005; Somel et al. 2013; Hughes et al. 2014). This relaxation is often explained by the fact that vision has become more important than smell in primates (Gilad et al. 2004). Note that having both DN/DS and PN/PS ratios around two suggest that non-synonymous mutations are neutral in these genes, which is consistent with the observation that many olfactory genes have become pseudogenes (Go and Niimura 2008).

The third highest scoring pathway is the *Beta-catenin phosphorylation cascade* pathway. Beta-catenin is involved in both cell adhesion and Wnt signaling (signal transduction). It also plays an important role during development and it can act as oncogene (Bienz 2005). This pathway is a good example of having an unusual combination of N/S coordinates, with PN/PS being much lower than DN/DS, while the separate ratios are not extreme outliers by themselves. This pattern is compatible with a recent strong purifying selection in humans resulting in very low PN/PS values (Figure 1D), possibly after an initial adaptive event.

Indeed, this is confirmed by the jackknife scores that show a constantly low PN/PS score across all genes. Two genes appear as outliers in opposite directions: the *APC* gene (a gene involved in tumor suppression and in synapse assembly, Rosenberg et al. 2008) plays a large role in the high DN/DS ratio of this gene set, while the *AXIN* gene has the opposite effect. When one or the other is removed the DN/DS ratio changes strongly, but when both are removed the DN/DS ratio (1.2) is close to the original value (1.25). We therefore posit that the signal shown by this pathway is indeed polygenic and not driven by the effect of a single gene.

The next high scoring gene set is *Glutathione conjugation*. This pathway contains many Glutathione S-Transferases (GSTs), which are involved in the protection of cells against oxidative stress and toxic foreign compounds (Hayes and Strange 2000). The jackknife scores of this pathway show that PN/PS ratios are typically high for all genes in this set, with some variation, whereas their DN/DS ratios is very stable and maintained at a moderately high level (Figure 1E). The fact that this pathway has an unusually high PN/PS ratio for shared SNPs could be due to some form of balancing selection. It suggests that the high diversity in GSTs is beneficial, possibly serving as a population wide protection against a large range of toxic factors in the environment. Indeed, GST polymorphisms have been related to drug sensitivity and are associated with disease susceptibility (Hayes and Strange 2000).

The *mRNA Splicing* pathway is a good candidate for being under extremely strong purifying selection as it shows particularly low values both for DN/DS and for PN/PS (Figure 1F). This pathway is an extreme example of a ‘housekeeping’ pathway, with genes expressed in all tissues. Earlier studies have indeed reported that housekeeping genes evolve more slowly than tissue specific genes, and that they are under stronger selective constraints (e.g. Zhang L and Li 2004).

The last two significant pathways, *Autoimmune thyroid disease* and *Graft-versus-host disease*, are related to immune response and have many genes in common. Their position relative to their expected distribution (Figure 1G-H), suggests that these sets could have been affected by positive selection or by a relaxation of selective constraints. However, both pathways contain many HLA genes (Table S1) that are normally considered as being under balancing selection (Solberg et al. 2008).Therefore long-term balancing selection would also explain this pattern.

### Impact of differential demography in African and non-African populations

Previous studies have shown that populations outside Africa have a higher proportion of deleterious mutations than those in Africa (Lohmueller et al. 2008; Subramanian 2012), compatible with the buildup of a mutation load since the expansion of modern humans out-of Africa (Peischl et al. 2013). We examined PN/PS ratios for SNPs that were either shared between Africans and non-Africans, or that were either private to Africans or private to non-Africans. In line with previous results (Lohmueller et al. 2008; Peischl et al. 2013), we find that the PN/PS ratio is larger for non-African specific SNPs than for SNPs private to Africans. In addition, shared SNPs show the lowest PN/PS ratio, and this for genes belonging or not to gene sets (Table 1). These results are consistent with the view that bottlenecks and range expansions of non-African populations have increased their PN/PS ratio relative to Africans (Peischl et al. 2013), and that purifying selection had more time to act on shared SNPs than on population specific SNPs, thus contributing to the elimination of a proportionally larger number of deleterious non-synonymous mutations.

In order to study the effect of potentially different demographic histories in Africans and non-Africans at the gene set level, we used our 2DNS test to compare the PN/PS ratios of private non-African SNPs and private African SNPs. As expected, PN/PS ratios are overall clearly larger for non-African SNPs than for African SNPs at the gene sets level (Figure 2, sign test: p<2.2e-16). In addition to showing the effect of past demography on the whole genome, our procedure reveals four significant outlier pathways (Table S2), which all have unusual N/S combinations given the genomic background. Among these, the *ALK1 signaling events* pathway is particularly interesting, since it has a markedly high non-African PN/PS ratio. This pathway plays a role in stimulating angiogenesis (Goumans et al. 2009), but also contains many genes (13 out of 23) that are involved in BMP signaling (Table S1). Interestingly, the *Signaling by BMP* pathway, which contains six genes in common with *ALK1 signaling events*, was found significant in our previous study contrasting levels of differentiation between continental groups at the gene set level (Daub et al. 2013). We then proposed that its role in iron metabolism and viral infection (Armitage et al. 2011) could be the key to the adaptive signal. Our present results confirm that this pathway might have been involved in local adaptation in non-African populations.

### Comparison between 2DNS and classical McDonald-Kreitman (MDK) tests

To illustrate the difference between our new approach and classical MDK tests, we performed an MDK test at the gene set level. In other words, we tested if some gene sets presented a DN/DS ratio larger than their PN/PS ratio. We inferred the significance of the results in two ways: with a conventional Fisher exact test on the 2×2 contingency table of gene set level DN, DS, PN and PS counts; and by comparing the gene set α value computed according to eq. (3) (Material and Methods) to an empirical null distribution built along the same principles as for the 2DNS test.

With a two sided Fisher exact test test (testing for deviation from DN/DS=PN/PS) we find 25 pathways scoring significant (q-value < 10%). All of them have a PN/PS > DN/DS, which would point towards purifying or balancing selection rather than to positive selection (Table S3). Interestingly, eight out of the ten highest scoring significant sets are directly related to immunity or response to pathogens. The first two pathways with DN/DS > PN/PS, indicative of positive selection, are *IL2-mediated signaling events* and *Beta-catenin phosphorylation cascade* and score (insignificant) q-values of 0.12 and 0.14 respectively. However, applying a one-sided Fisher exact test, where we explicitly try to detect cases where DN/DS > PN/PS, produces no significant pathways at FDR level 10%. (Table S4). We note however that a QQ plot analysis of the p-values of both Fisher exact tests shows clear systematic departures from expectations. Indeed, the p-values are under- and overestimated for the two sided and for the one sided Fisher exact test, respectively (Figure S6), implying that the two sided test is too liberal, and the one-sided test is too conservative.

The comparison of gene set *α* values to an empirical null distribution results in several high scoring pathways, but none of them are significant after correcting for multiple tests (Table S5), suggesting that a test based on *α* values is less powerful than our 2DNS test. Moreover, we notice that most of the top-ranking pathways (shown as dark red points on Figure S7) have a high *α* value associated with a low DN/DS and even lower PN/PS. It suggests that positive selection is more easily detected in (or is more likely to act on) slowly evolving genes.

## Discussion

Our 2DNS test has several advantages over classical methods such as the McDonald-Kreitman test. First of all, we focus on the detection of selection in functional groups of genes instead of single genes. Not only is a biological pathway or gene network a more natural unit to test for selection, but by pooling genes belonging to the same gene set we avoid the exclusion of many genes that have undefined N/S ratios. Second, our test allows one to detect different selective regimes, whereas classical tests are often designed to evidence only one type of selection, usually positive selection. As proposed in Figure 3, we can infer which selective regime could have acted on an outlier gene set from its position in the PN/PS-DN/DS plane. Still, different selective processes can yield similar patterns leading to ambivalent interpretations. However, we see this as a problem of the underlying biology (different processes generate similar patterns) rather than of the 2DNS test *per se*. In these cases one could inspect the function of the genes of a candidate pathway in more detail to gain insight on the type of selection that might have acted on the gene set.

**Figure 3.**
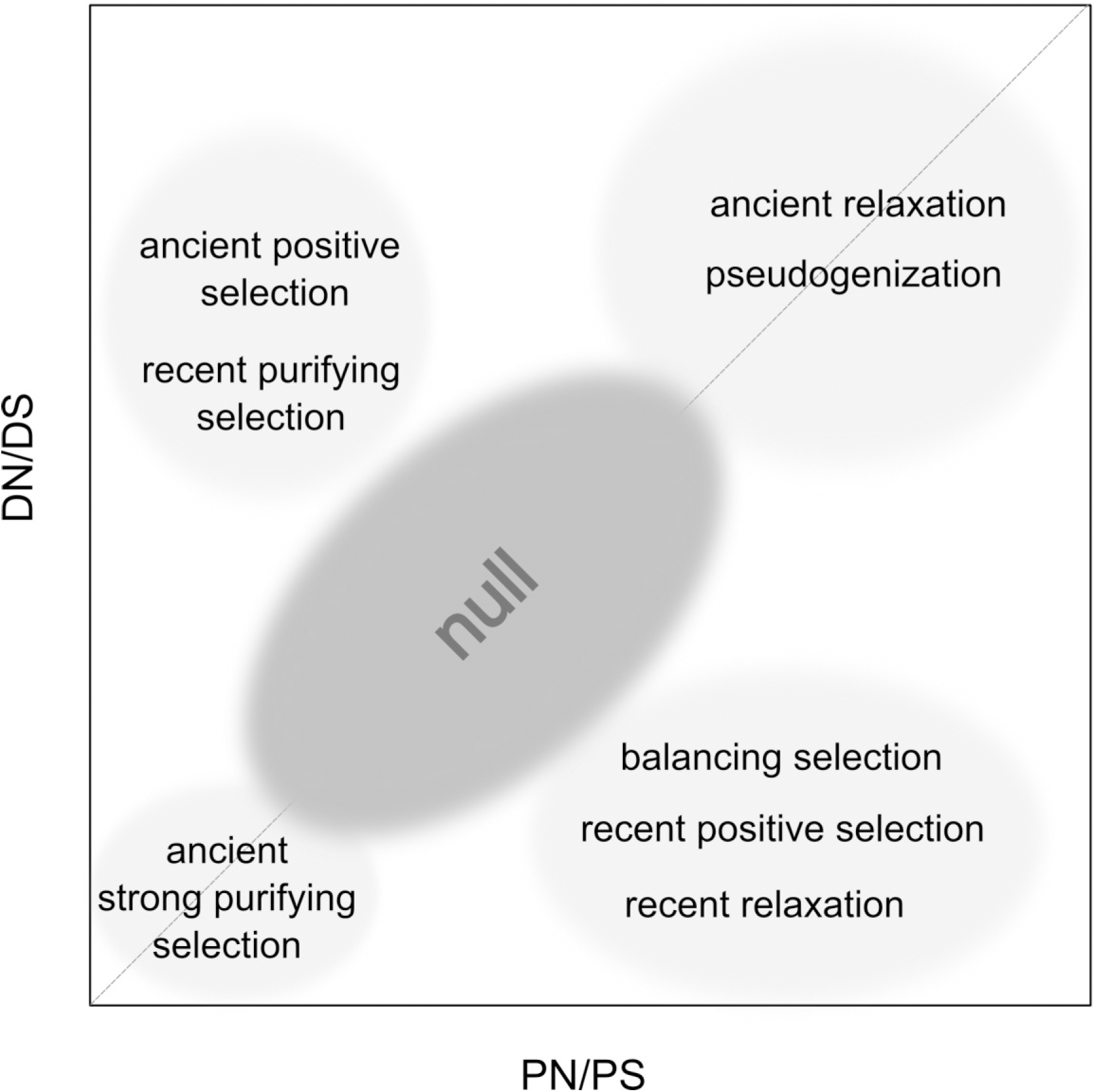
Potential evolutionary forces having acted on significant outlier gene sets depending on their position in the N/S plane.

Our 2DNS test is similar in spirit to the 2D neutrality test proposed by Innan (2006), which incorporates two summary statistics to test genes for signals of selection. However, the statistics used by Innan are indirect measures of neutrality, and the interpretation of these pairs of values is not straightforward. Rather, instead of combining two selection tests, we decompose here the MDK test into its two constituent factors, DN/DS and PN/PS, which allows us to gain more insight into underlying evolutionary pressures.

A third important advantage of our method is that it compares gene sets against a genome-wide empirical null distribution, and thus controls for potential demographic events (similar to e.g. Nielsen et al. 2005) and genome-wide selective forces, most notably background selection (e.g. background selection, Hernandez et al. 2011). It has indeed been noticed that large genomic data sets contain their own null distribution (Efron 2013), implying they can be used to evidence outliers. Note however that our approach does not test against any pre-specified evolutionary model, and it is therefore not a test of neutrality since gene sets found to be non-significant by our approach can be under selection. With our 2DNS test, we are thus trying to find extreme outliers, such as gene sets that have atypical patterns of diversity compared to other gene sets, which makes it a rather conservative method, as we build a null distribution that closely reflects the properties of the gene sets themselves. However, we have shown that using a naïve null distribution, for example by simply randomly sampling from all genes, would yield many false positives (Figure S4), because this null distribution would not reflect the inherent properties of gene sets in general. This is especially true because pathways described in current databases are probably a biased subset of existing biological pathways.

In line with earlier studies (Bustamante et al. 2005; Chimpanzee Sequencing and Analysis Consortium 2005; Zhang L and Li 2005; Eyre-Walker and Keightley 2009; Messer and Petrov 2013), we find more evidence for purifying rather than positive selection, reflected for example in lower DN/DS ratios than PN/PS ratios, both at the gene and gene set level (Table 1). However, a possible explanation for finding only a few examples of positive selection is that it does not affect mostly coding regions, but rather regulatory regions or other functional parts of the genome, leading to variation in gene expression or epigenetic differences (Ponting and Lunter 2006; Enard et al. 2014). The 2DNS test could be adapted to study such categories of genomic data, for example comparing mutations in transcription factor binding sites to putative neutral flanking sites, as other metrics than synonymous or non-synonymous state can be used to assign the level of functional or selective constraints; e.g., site conservation scores (e.g. GERP scores, Cooper et al. 2005).

Our 2DNS test is not restricted to finding gene sets with unusual DN/DS-PN/PS pairs in the human branch and can easily be extended to other species, or applied to compare evolutionary rates between species. The test is also suitable to compare the diversity levels among polymorphisms between groups with a different demographic history, as exemplified in Figure 3, where we compare SNPs private to African and to non-African populations.

In the NS plane many small gene sets show very unusual joint N/S ratios (Figure 1A), but they are not found significant because their associated null distributions are very wide (e.g. Figure 1I). This shows that one should be cautious with groups of genes that have high N/S ratios, as such values can arise by chance alone, but it also suggests that our test might lack power to detect outlier selection regimes in small gene sets. In addition, it is likely that selection has not affected whole pathways but only some sub-sets, and that this limited signal might be difficult to detect at the whole pathway level. Alternatively, different evolutionary forces might also act on distinct subsets of genes, and their signal could cancel each other’ s out when examining whole pathways, leading to a reduced power of our approach. Note that this problem also occurs when testing single genes, since different exons, introns or regulatory regions can be affected by different selective forces. It would thus be useful to extend our test to detect selection in pathway sub-modules, which should be the object of future work.

Our previous attempt at detecting selection at the gene set level within human populations revealed several pathways involved in immune response to be potentially under positive selection (Daub et al. 2013). Interestingly, our present study does not show strong signals of positive selection, but rather of overall purifying selection. These differences can be due to several reasons. First, our previous study used a statistic (hierarchical *F*_*ST*_) detecting differences between continental groups that should have emerged recently, whereas our 2DNS statistics are more sensitive to older events. We contrast old mutations that have occurred on the human lineage to polymorphisms shared between Africans and non-Africans (Figure 1), and which should thus have occurred in the ancestral human populations prior to the exit out of Africa. Note however that when we contrast patterns of polymorphism between Africans and non-Africans for younger mutations (Figure 2), we indeed find one pathway (*ALK1 signaling events*) showing signs of positive selection in non-Africans, and that this pathway share several genes with a pathway that we previously identified as being under positive selection (*signaling by BMP*). Second, our current testing procedure is more stringent than that used previously. We are indeed looking for outliers in a null distribution that takes into account selective constraints acting on pathways as well as past demographic effects, whereas we previously based our test on a null model of human evolution only taking into account global levels of differentiation between human populations. Third, we focused here strictly on coding regions, whereas we also considered neighboring regions in our previous study, thus including regulatory and enhancer regions that have been recently shown to bear the strongest signals of positive selection in humans (Enard et al. 2014). These differences in methodology and in the investigated time scales could thus explain the apparent discrepancies found between these two studies. This also indicates that different forces have acted on pathways at different periods of human evolution.

## Acknowledgements

We thank Adam Eyre-Walker for his critical comments on a previous version of the manuscript. This work was supported by the Swiss National Science Foundation (grant number PDFMP3-130309 to LE).

## Text S1

### Estimation of variance component among gene sets

To investigate whether gene-level DN/DS (and PN/PS) ratios are more similar within than between gene sets, we estimated the variance component attributable to differences among gene sets(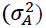) under a classical analysis of variance (ANOVA) framework (see e.g. Sokal and Rohlf 1981, p. 216) where a given N/S ratio *(x*_*ij*_) in gene *j* of gene set *i* is modelled as

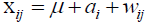

where *μ* is the unknown expectation of x_*ij*_, *a*_*i*_ is an effect due to the gene set i, and *w*_*ij*_ is the effect due to gene *j*. The total variance in N/S ratio 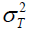 is estimated as 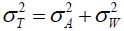, where 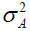 and 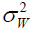 are the variance components due to differences between gene sets and between genes within gene sets, respectively.

Assuming that we have *N* genes and *S* gene sets, we can construct a standard ANOVA table

| Source of variation | Degrees of freedom | Mean squares (MS) | Expected MS |
| --- | --- | --- | --- |
| Among gene sets | $df_A = S - 1$ | $MS_A = SS_A/df_A$ | $\sigma_W^2 + n_0 \sigma_A^2$ |
| Within gene sets | $df_W = N - S$ | $MS_W = SS_W/df_W$ | $\sigma_W^2$ |

where the sums of squares are obtained as 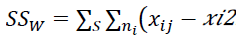.

The average gene set size, *n*_*0*_, is obtained as 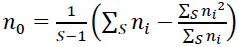 where *n*_*i*_ is the size of gene set *i*.

The variance components are estimated as ***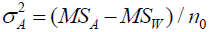*** and ***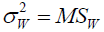***.

**Table S1.** Genes in top scoring gene sets (separate excel file TableS1.xlsx). For each gene in the gene sets the DN and DS counts and PN and PS counts for the SNP groups are reported. For the *ALK1 Signaling* pathway the involvement in BMP signaling is indicated (having a GO term or being part of a pathway related to BMP signaling), as inferred from the NCBI Entrez Gene database (Maglott, et al. 2011).

**Table S2.**
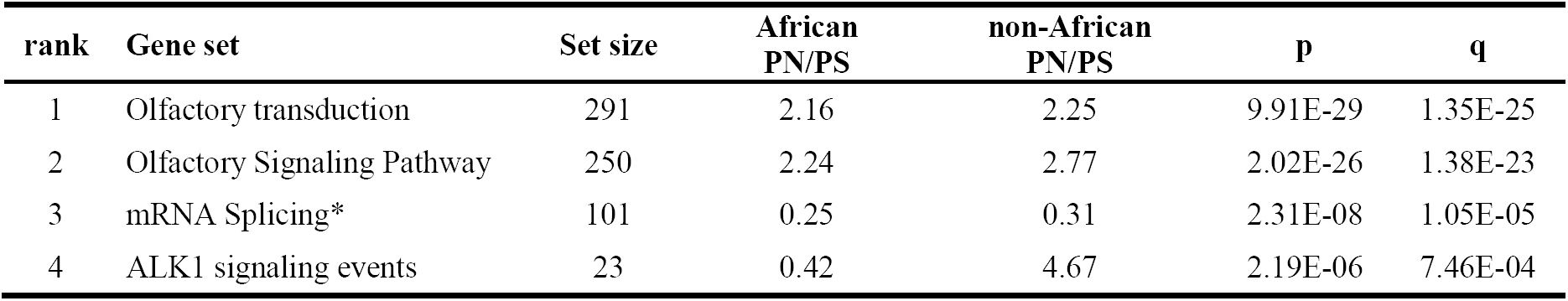
Pathways scoring a q-value < 0.1 in the 2DNS test, applied on polymorphisms unique to African populations (African PN/PS) and those unique to non-African populations (non-African PN/PS).

**Table S3.**
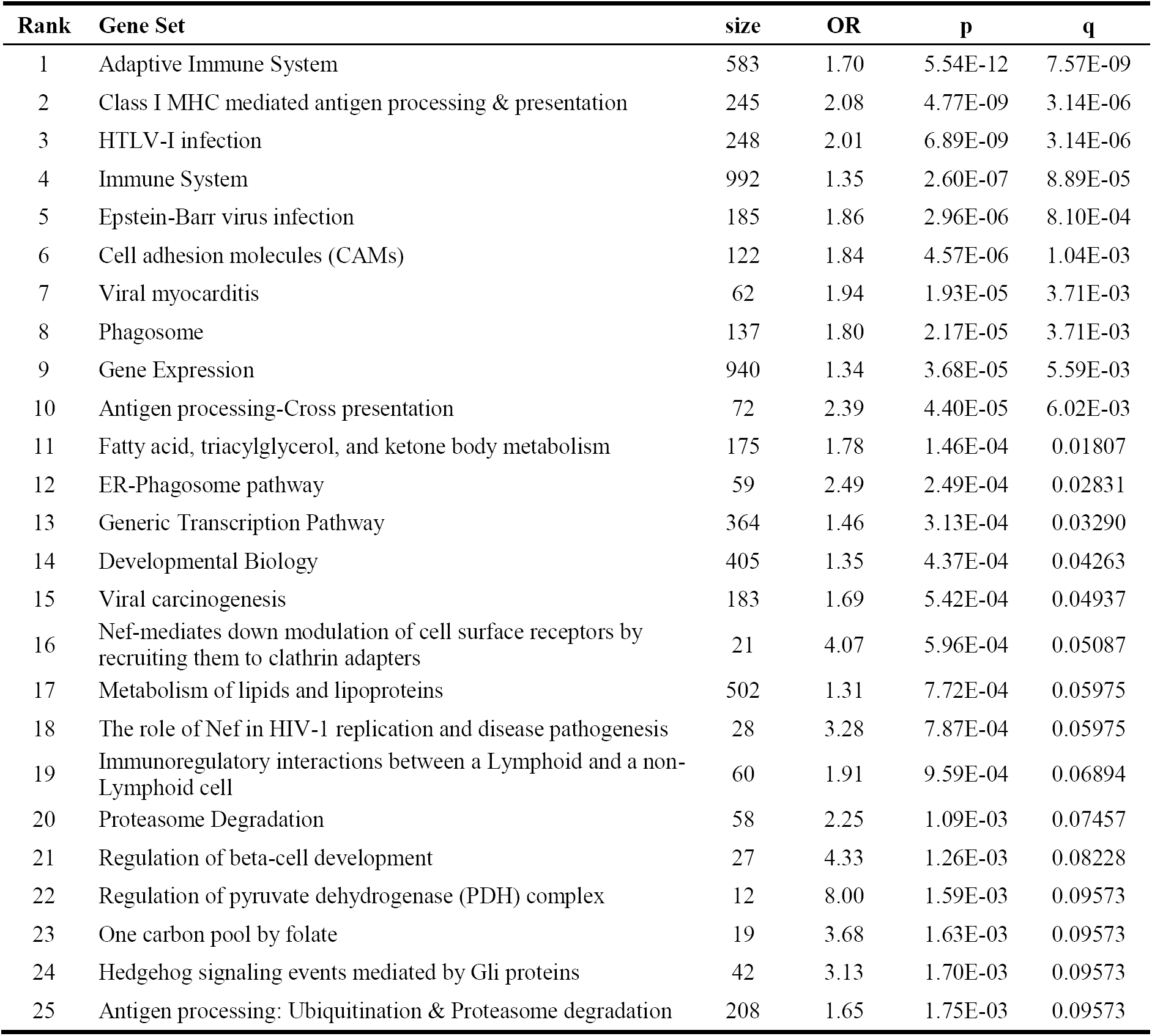
Results from classical McDonald-Kreitman test, comparing shared human SNPs with fixed mutations in the human branch. Here significance of a gene set is inferred with a two-sided Fisher exact test, taking as null-hypothesis that the odds ratio of the contingency table OR=(DS·PN)/(DN·PS)=1. Gene sets scoring a q-value<0.1 are reported.

| Rank | Gene Set | size | OR | p | q |
| --- | --- | --- | --- | --- | --- |
| 1 | Adaptive Immune System | 583 | 1.70 | 5.54E-12 | 7.57E-09 |
| 2 | Class I MHC mediated antigen processing & presentation | 245 | 2.08 | 4.77E-09 | 3.14E-06 |
| 3 | HTLV-I infection | 248 | 2.01 | 6.89E-09 | 3.14E-06 |
| 4 | Immune System | 992 | 1.35 | 2.60E-07 | 8.89E-05 |
| 5 | Epstein-Barr virus infection | 185 | 1.86 | 2.96E-06 | 8.10E-04 |
| 6 | Cell adhesion molecules (CAMs) | 122 | 1.84 | 4.57E-06 | 1.04E-03 |
| 7 | Viral myocarditis | 62 | 1.94 | 1.93E-05 | 3.71E-03 |
| 8 | Phagosome | 137 | 1.80 | 2.17E-05 | 3.71E-03 |
| 9 | Gene Expression | 940 | 1.34 | 3.68E-05 | 5.59E-03 |
| 10 | Antigen processing-Cross presentation | 72 | 2.39 | 4.40E-05 | 6.02E-03 |
| 11 | Fatty acid, triacylglycerol, and ketone body metabolism | 175 | 1.78 | 1.46E-04 | 0.01807 |
| 12 | ER-Phagosome pathway | 59 | 2.49 | 2.49E-04 | 0.02831 |
| 13 | Generic Transcription Pathway | 364 | 1.46 | 3.13E-04 | 0.03290 |
| 14 | Developmental Biology | 405 | 1.35 | 4.37E-04 | 0.04263 |
| 15 | Viral carcinogenesis | 183 | 1.69 | 5.42E-04 | 0.04937 |
| 16 | Nef-mediates down modulation of cell surface receptors by recruiting them to clathrin adapters | 21 | 4.07 | 5.96E-04 | 0.05087 |
| 17 | Metabolism of lipids and lipoproteins | 502 | 1.31 | 7.72E-04 | 0.05975 |
| 18 | The role of Nef in HIV-1 replication and disease pathogenesis | 28 | 3.28 | 7.87E-04 | 0.05975 |
| 19 | Immunoregulatory interactions between a Lymphoid and a non-Lymphoid cell | 60 | 1.91 | 9.59E-04 | 0.06894 |
| 20 | Proteasome Degradation | 58 | 2.25 | 1.09E-03 | 0.07457 |
| 21 | Regulation of beta-cell development | 27 | 4.33 | 1.26E-03 | 0.08228 |
| 22 | Regulation of pyruvate dehydrogenase (PDH) complex | 12 | 8.00 | 1.59E-03 | 0.09573 |
| 23 | One carbon pool by folate | 19 | 3.68 | 1.63E-03 | 0.09573 |
| 24 | Hedgehog signaling events mediated by Gli proteins | 42 | 3.13 | 1.70E-03 | 0.09573 |
| 25 | Antigen processing: Ubiquitination & Proteasome degradation | 208 | 1.65 | 1.75E-03 | 0.09573 |

**Table S4.** Results from classical McDonald-Kreitman test, comparing shared human SNPs with fixed mutations in the human branch. Here we tested the significance of a gene set with a one-sided Fisher exact test, taking as null-hypothesis that the odds ratio of the contingency table OR=(DS·PN)/(DN·PS)>=1. The 10 highest scoring gene sets are shown.

| Rank | Gene Set | size | OR | p | q |
| --- | --- | --- | --- | --- | --- |
| 1 | IL2-mediated signaling events | 50 | 0.29 | 0.0017 | 1 |
| 2 | Beta-catenin phosphorylation cascade | 17 | 0.07 | 0.0026 | 1 |
| 3 | IL-2 Signaling Pathway | 75 | 0.44 | 0.0052 | 1 |
| 4 | Transport of inorganic cations/anions and amino acids/oligopeptides | 94 | 0.62 | 0.0067 | 1 |
| 5 | Abortive elongation of HIV-1 transcript in the absence of Tat | 22 | 0.10 | 0.0073 | 1 |
| 6 | Salivary secretion | 83 | 0.66 | 0.0082 | 1 |
| 7 | Regulation of Insulin-like Growth Factor (IGF) Activity by Insulin-like Growth Factor Binding Proteins (IGFBPs) | 17 | 0.32 | 0.0103 | 1 |
| 8 | HIV-1 elongation arrest and recovery* | 30 | 0.26 | 0.0106 | 1 |
| 9 | Pancreatic secretion | 94 | 0.65 | 0.0108 | 1 |
| 10 | Regulation of AMPK activity via LKB1 | 14 | 0.14 | 0.0128 | 1 |

**Table S5.** Results from the alpha test, where α-values (1-PN·DS/PS·DN) of gene sets were contrasted against a null distribution of random sets. Here we used shared human SNPs and fixed mutations in the human branch. The 10 highest scoring gene sets are shown.

| Rank | Gene Set | size | $\alpha$ | p | q |
| --- | --- | --- | --- | --- | --- |
| 1 | mRNA Splicing - Minor Pathway | 39 | 0.84 | 0.00074 | 0.443 |
| 2 | IL2-mediated signaling events | 50 | 0.71 | 0.00078 | 0.443 |
| 3 | Beta-catenin phosphorylation cascade | 17 | 0.93 | 0.00108 | 0.443 |
| 4 | C-MYC pathway | 21 | 1.00 | 0.00130 | 0.443 |
| 5 | Processing of Capped Intronless Pre-mRNA | 20 | 1.00 | 0.00348 | 0.750 |
| 6 | Canonical Wnt signaling pathway | 29 | 0.70 | 0.00383 | 0.750 |
| 7 | Serotonin Receptor 4/6/7 and NR3C Signaling | 19 | 0.80 | 0.00458 | 0.750 |
| 8 | IL-2 Signaling Pathway | 75 | 0.56 | 0.00494 | 0.750 |
| 9 | Platelet calcium homeostasis | 19 | 0.72 | 0.00559 | 0.750 |
| 10 | Glucagon signaling in metabolic regulation | 31 | 0.68 | 0.00595 | 0.750 |

**Figure S1.**
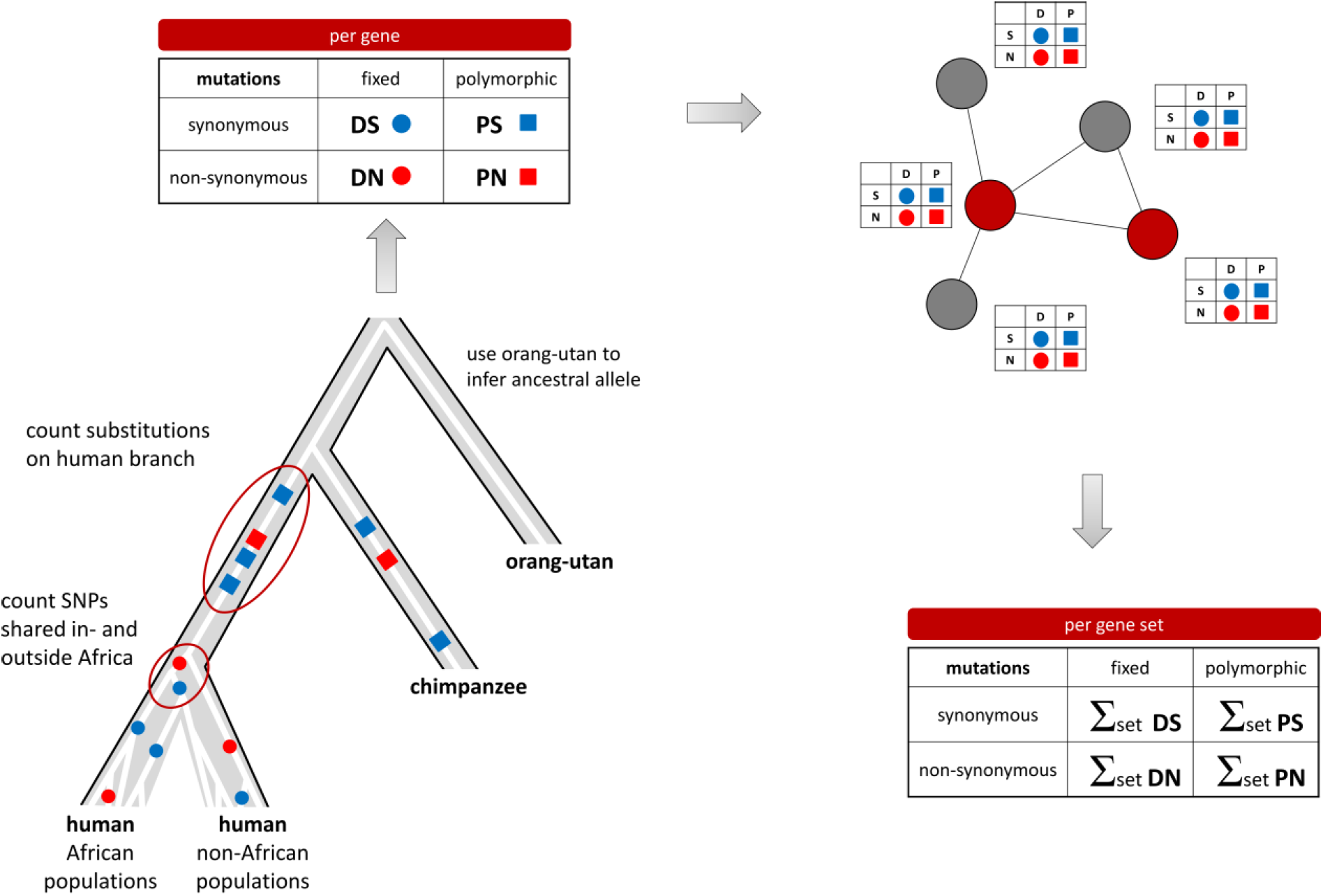
Overview of the pipeline used to generate gene set level DN/DS and PN/PS ratios.

**Figure S2.**
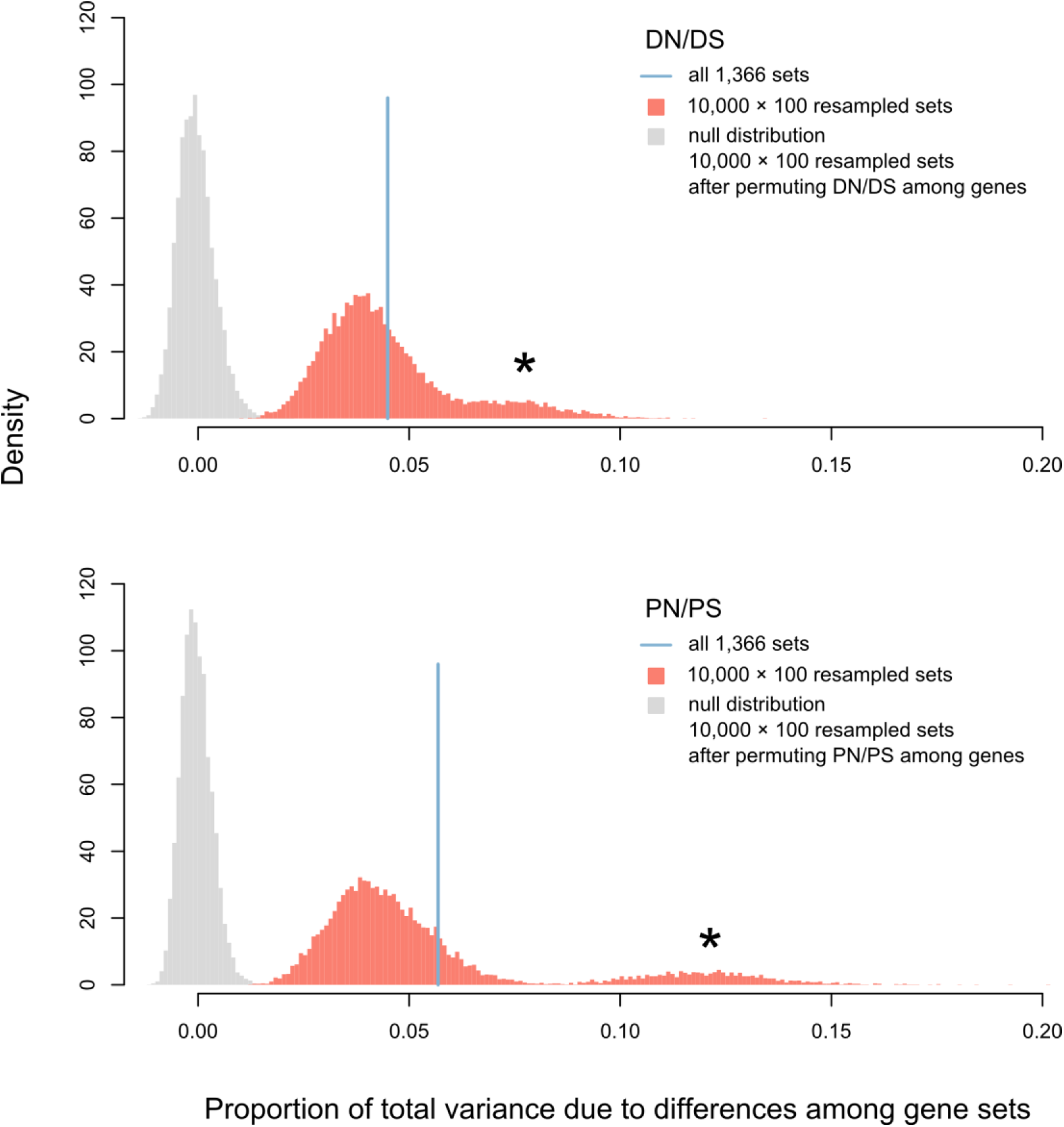
Proportion of the total variance of gene level DN/DS and PN/PS ratios explained by differences among gene sets. We repeatedly (*N*=10,000) sampled randomly 100 sets from the whole list of 1,366 gene sets and computed the proportion of the total variance due to differences among these 100 gene sets. A null distribution was created following the same procedure, but with each repetition the DN/DS (or PN/PS) ratios were permuted among genes (grey). We show here that both for DN/DS ratios as for PN/PS ratios, the distribution of the variance component due to differences between gene sets is significantly larger than zero (null distribution). This means that gene sets differ significantly from each other, implying that genes that are part of the same set are significantly more similar than by chance alone for their N/S ratios. It suggests that genes that are part of the same set tend to share the same evolutionary regime. Note that the second peak at higher values (indicated with a *) is caused by the two highest scoring gene sets in the 2DNS test, the *Olfactory Signaling Pathway* and *Olfactory Transduction*. The variance component computed on all 1,366 sets is indicated with a blue line. Genes with undefined N/S ratios, having DS=0 (A) or PS=0 (B), were excluded.

**Figure S3.**
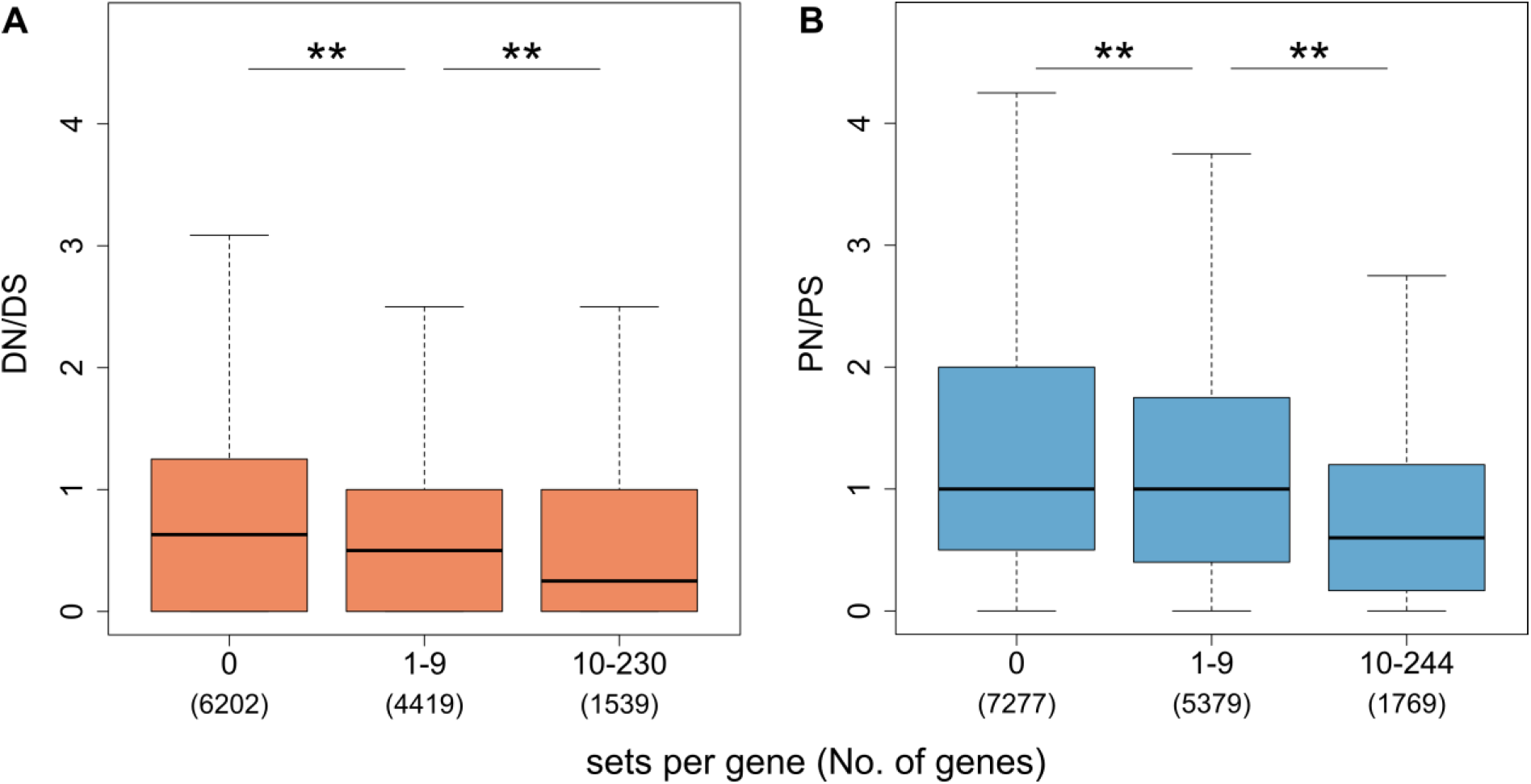
Genes that are part of many pathways are more conserved. Distribution of DN/DS ratios (A) and PN/PS ratios (B) of genes that are part of 0, 1-9 or >9 pathways. The number of genes in each category are shown in parentheses. Genes with undefined N/S ratios, having DS=0 (A) or PS=0 (B), were excluded. Significant differences between the groups as inferred with a Mann-Whitney test are marked with a ** (p-value < 1e-6).

**Figure S4.**
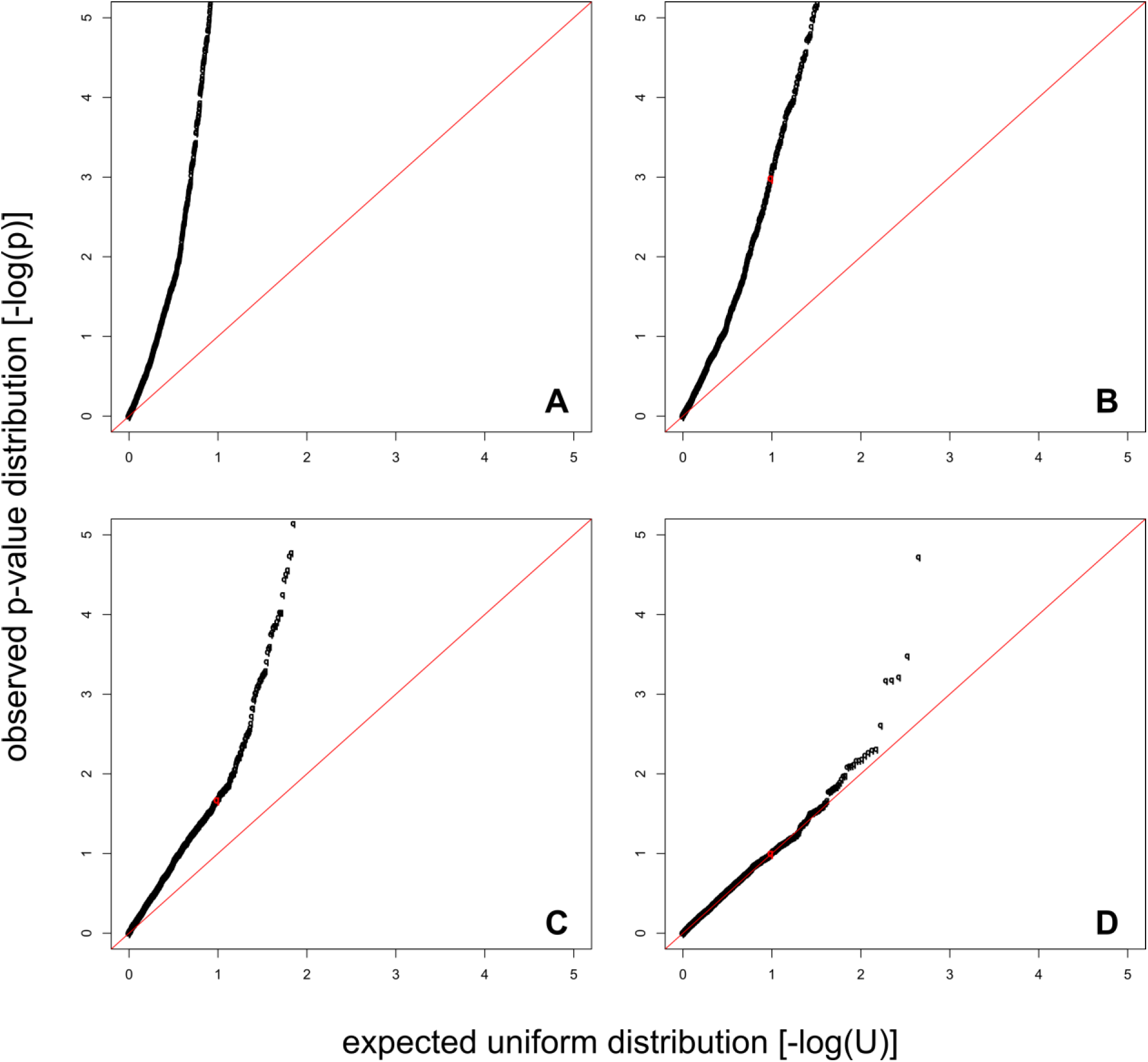
QQ plots of the distribution of log transformed p-values from the 2D test against a log transformed uniform distribution of values between 0 and 1. Here the 2D test compared DN/DS ratios of fixed mutations on the human branch against PN/PS ratios of shared human polymorphisms. The null distribution is build according to the following sampling schemes: (A) randomly sampling from the whole gene list G; (B) sampling from all genes that are part of at least one gene set; (C) sampling genes with a probability proportional to the number of pathways a gene occurs in; and (D) sampling from unions of similar gene sets (the empirical null distribution taking into account gene set properties). All null distributions were created for specific set length bins and pathways were tested against the corresponding null distribution. The red dot marks the 90% quantile of the p-value distribution.

**Figure S5.**
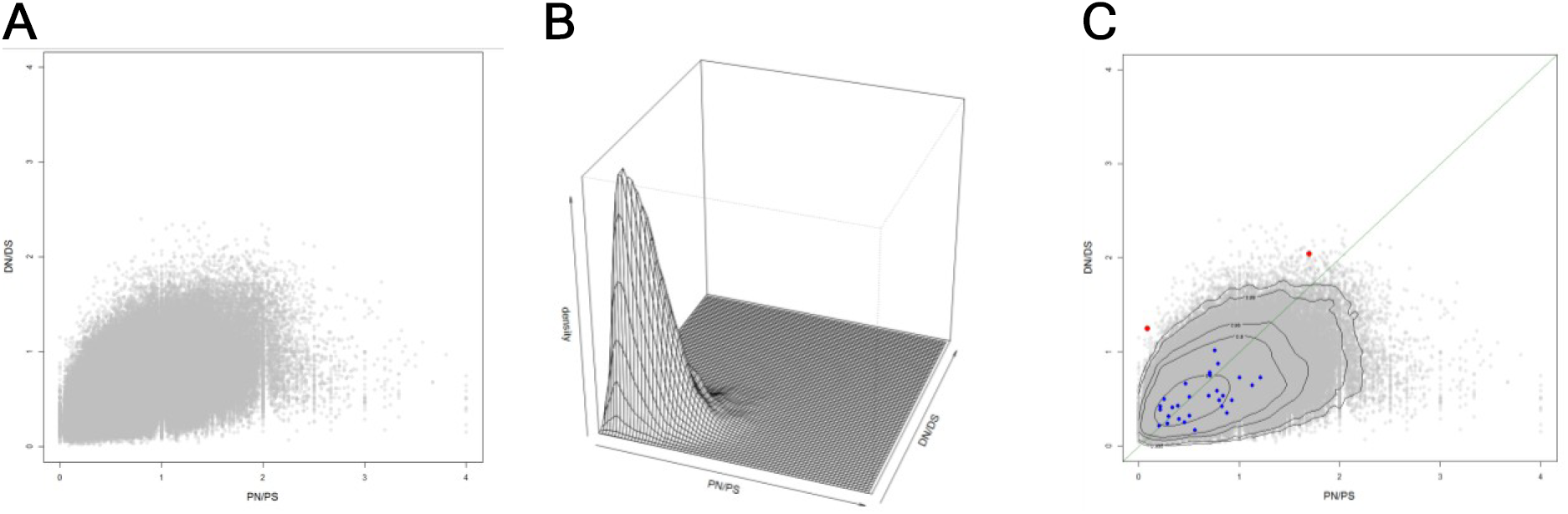
After generating an empirical null distribution of random gene sets (A), the joint density of the null distribution is estimated (B), and the p-values of the gene sets are estimated (C) for a given set length bin (blue dots: non-significant sets, red dots: significant sets).

**Figure S6.**
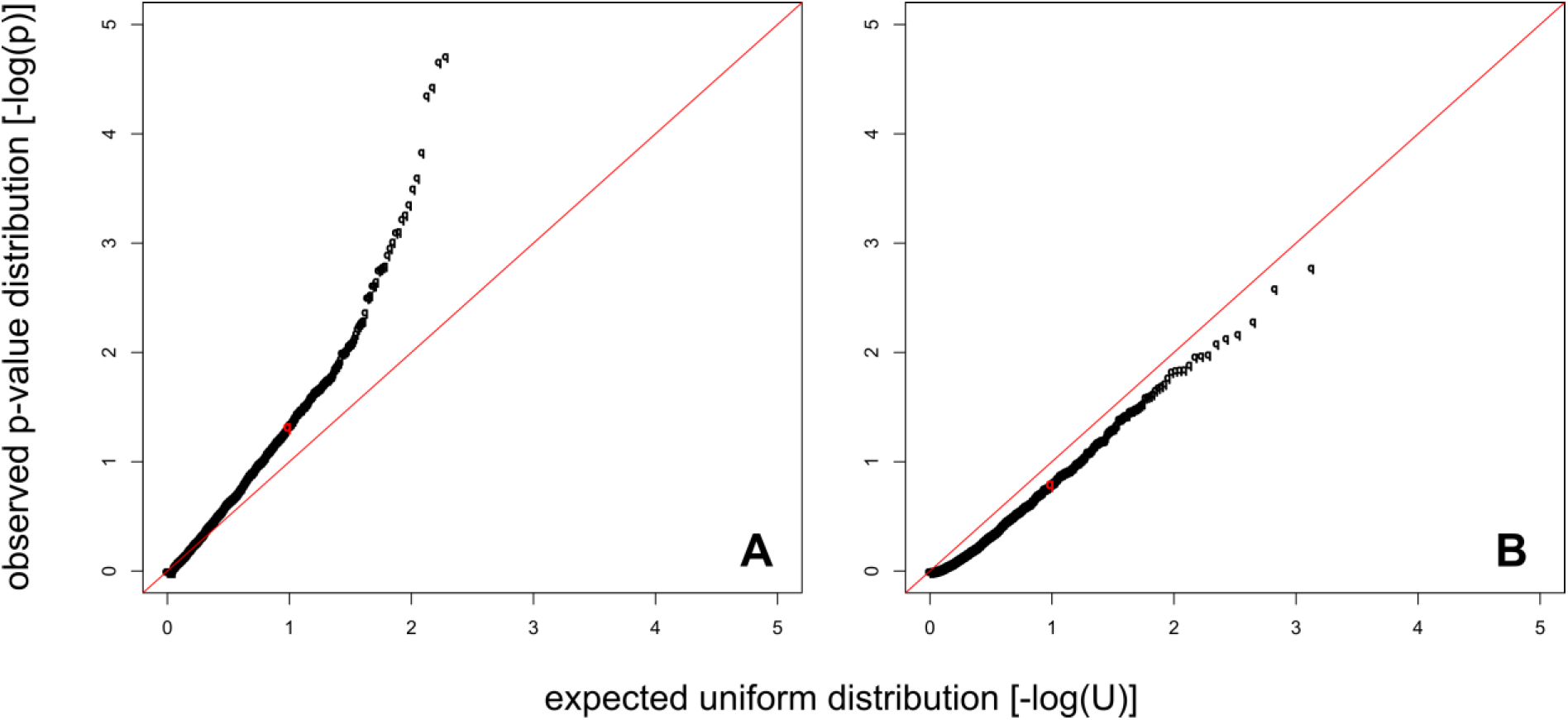
QQ plot of the distribution of log transformed p-values from a gene set level MDK test against a log transformed uniform distribution of values between 0 and 1. Here the test compared shared human SNPs with fixed mutations in the human branch. Significance of a gene set is inferred with (A) a **two-sided** Fisher exact test, taking as null-hypothesis that the odds ratio of the contingency table OR=(DS·PN)/(DN·PS)=1, or with (B) a **one-sided** Fisher exact test, taking as null-hypothesis that the odds ratio of the contingency table OR=(DS·PN)/(DN·PS)≤1. The red dot marks the 90% quantile of the p-value distribution.

**Figure S7.**
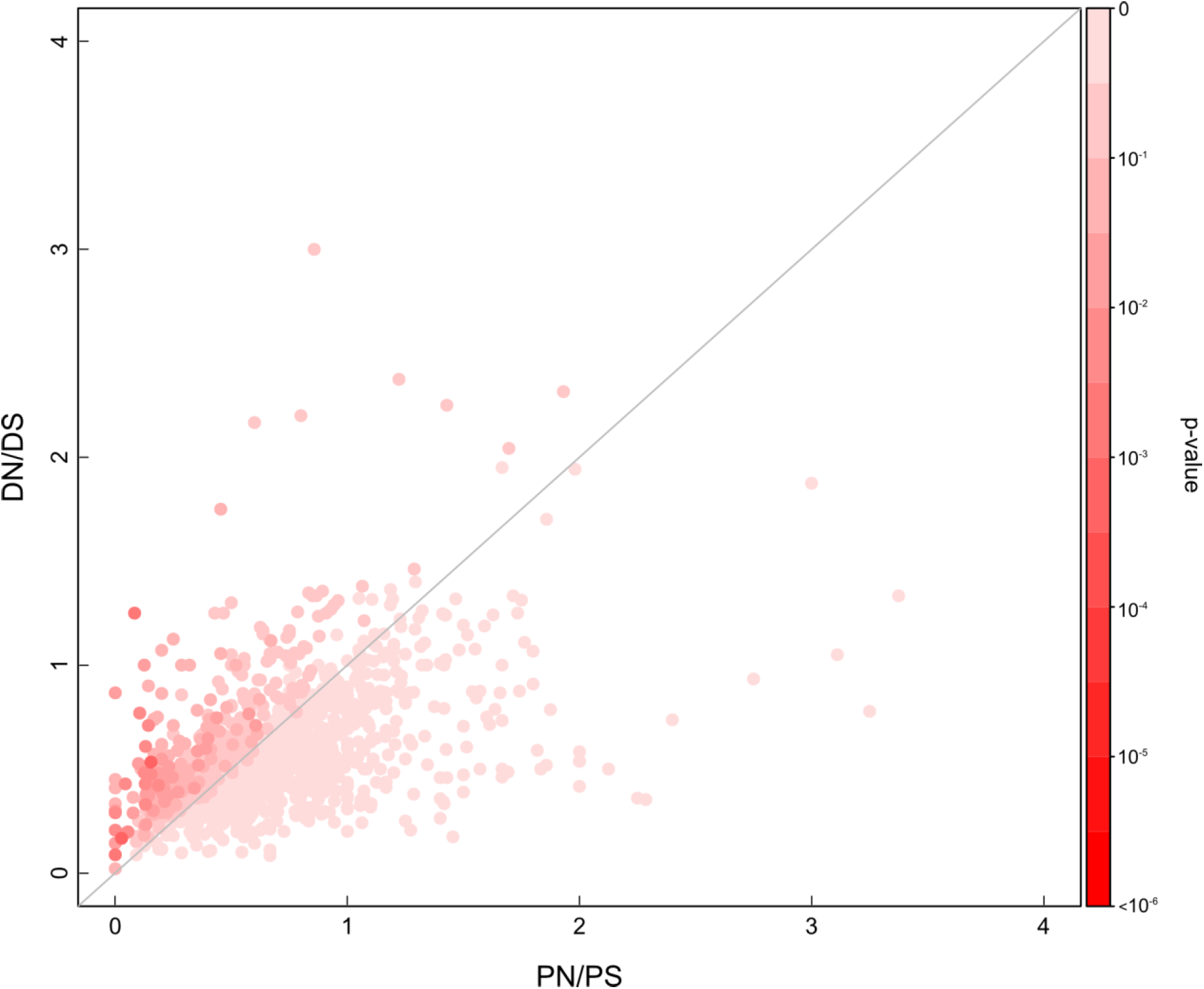
Results of the McDonald-Kreitman α test for polygenic selection, comparing DN/DS and PN/PS ratios for all tested pathways. Here, shared SNPs (SNPs that are polymorphic both in African and non-African populations) are compared to substitutions in the human branch. Each dot represents a pathway with color corresponding to its significance. α-values (computed as 1-PN·DS/PS·DN) of gene sets were compared to an empirical null distribution of random sets to infer p-values. No pathway scored significant at a FDR level of 10% (q-value < 0.1).

